# Scalable spatial single-cell transcriptomics and translatomics in 3D thick tissue blocks

**DOI:** 10.1101/2024.08.05.606553

**Authors:** Xin Sui, Jennifer A. Lo, Shuchen Luo, Yichun He, Zefang Tang, Zuwan Lin, Yiming Zhou, Wendy Xueyi Wang, Jia Liu, Xiao Wang

**Author notes:** These authors contributed equally (Xin Sui, Jennifer A. Lo).

## Abstract

Characterizing the transcriptional and translational gene expression patterns at the single-cell level within their three-dimensional (3D) tissue context is essential for revealing how genes shape tissue structure and function in health and disease. However, most existing spatial profiling techniques are limited to 5-20 µm thin tissue sections. Here, we developed Deep-STARmap and Deep-RIBOmap, which enable 3D *in situ* quantification of thousands of gene transcripts and their corresponding translation activities, respectively, within 200-µm thick tissue blocks. This is achieved through scalable probe synthesis, hydrogel embedding with efficient probe anchoring, and robust cDNA crosslinking. We first utilized Deep-STARmap in combination with multicolor fluorescent protein imaging for simultaneous molecular cell typing and 3D neuron morphology tracing in the mouse brain. We also demonstrate that 3D spatial profiling facilitates comprehensive and quantitative analysis of tumor-immune interactions in human skin cancer.

## Introduction

The spatial regulation of gene expression and translation is critical for tissue function^1–6^. *In situ* profiling technologies enable the study of both the transcriptome and translatome within their original spatial contexts^7–10^. However, most spatial omics techniques are confined to analyzing thin tissue sections (5-20 µm). Many functional and anatomical studies in tissue biology require 3D profiling in tissue blocks across multiple cellular layers^11,12^. For instance, in neuroscience, 3D morphological profiling and long-range projection mapping^13–16^, *in situ* electrophysiology^17–22^, and *in vivo* neural activity imaging^23–25^ in the brain require direct measurements in 3D brain volumes (100-300 µm) where thin tissue sections are inadequate. In cancer pathology, 3D samples offer a more accurate representation of tumor architecture, microenvironment, and cell-cell interactions in patient samples^26^.

Although reconstructing 3D volumes using serial thin sections is feasible, this method faces three significant challenges. Firstly, tissue sectioning often fragments cells, resulting in partial RNA readouts and an increased risk of RNA content loss during handling, complicating the accurate analysis of the transcriptome. Secondly, tissue deformation during sectioning presents a persistent challenge for computational reconstruction methods. Thirdly, this approach requires substantial manual labor. Therefore, developing effective spatial omics methods for 3D profiling from thick tissue blocks is imperative.

Current thick-tissue spatial profiling approaches achieved quantitative *in situ* measurements of transcriptome in thick samples using single or multi-round Fluorescence *In Situ* Hybridization (FISH)^27–40^, but are limited in several aspects. The number of genes they can analyze, typically fewer than 300, and the size of the imaging areas, often restricted to a single brain region. These limitations arise primarily because these methods often employ linear coding instead of exponential coding, and rely on RNA integrity to preserve the imaging signal, significantly reducing their efficiency of RNA detection, scalability of gene numbers and tissue volume, and flexibility of sample handling and imaging time^28–40^. Additionally, the displacement of RNA molecules between imaging rounds further restricts the number of imaging cycles that can be performed^27^. Moreover, current thick-tissue spatial profiling methods are limited to mapping spatial transcriptomics and lack the capability to map the translatome, thus hindering multiplexed characterization of gene translation at single-cell resolution.

Here, we have developed Deep-STARmap and Deep-RIBOmap to address the aforementioned limitations by introducing a novel and scalable strategy for probe synthesis and embedding as well as robust cDNA amplicons crosslinking, enabling scalable *in situ* quantification of thousands of RNA transcripts and their respective translational activities within large intact thick tissue samples. Utilizing Deep-STARmap and Deep-RIBOmap, we profiled the transcription and translation of 1,017 genes in intact mouse brain tissue at 300 nm voxel size within a thick hydrogel-tissue scaffold, revealing heterogeneity in protein translation across cell types. Additionally, by combining our method with multicolor fluorescence labeling (Tetbow)^14^, we simultaneously profiled neuronal morphology and molecular signatures in single cells, achieving multimodal mapping of the adult mouse brain in a scalable manner. Lastly, we demonstrated the applicability of our method on human cutaneous squamous cell carcinomas (cSCC) samples, uncovering tumor-immune interactions with more accurate and quantitative spatial distributions compared to thin tissue analyses. We anticipate that Deep-STARmap and Deep-RIBOmap will also yield important biological insights into the pathophysiology of cancers and other diseases.

## Results

### Deep-STARmap and Deep-RIBOmap workflow

We designed the workflow of Deep-STARmap and Deep-RIBOmap (**Fig. 1a**) as follows: it begins with the hybridization of pre-designed oligonucleotide probe sets to target either all RNA molecules of a gene or ribosome-bound RNAs, respectively, in PFA-fixed tissues followed by hydrogel matrix embedding; the samples are then subjected to protein digestion and lipid removal to enhance enzyme penetration, ensuring sufficient depth coverage in thick tissue samples; subsequently, *in situ* cDNA amplicons are synthesized through enzymatic ligation and rolling circle amplification (RCA); each cDNA amplicon contains a pre-designed gene-specific identifier, which is finally decoded through cyclic sequencing, imaging, and stripping steps (SEDAL sequencing^30^). In comparison with previously published thick-tissue STARmap protocol (linear encoding, 28 genes), the new developments of Deep-STARmap and Deep-RIBOmap solved the issues of scalable probe preparation, cDNA amplicon anchoring, signal decay, and translation mapping capability as detailed below.

**Fig. 1.**
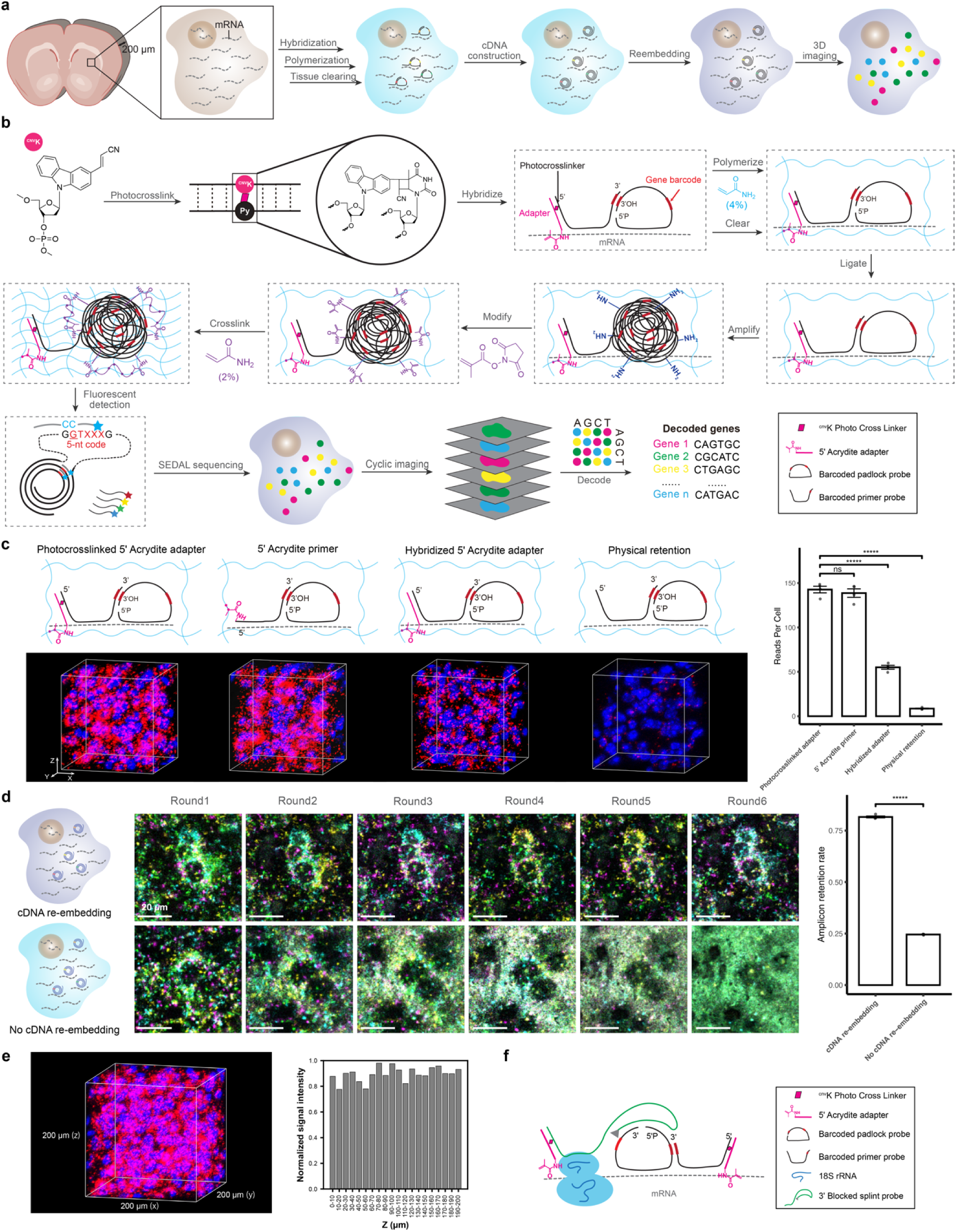
Deep-STARmap and Deep-RIBOmap enable spatiotemporally resolved transcriptomics and translatomics in 200 µm thick tissue blocks. **a**, Schematic summary of Deep-STARmap and Deep-RIBOmap workflow. **b**, *In situ* sequencing of transcriptional states in thick tissue blocks: The primer, featuring a flanking linker sequence at its 5′ end, is covalently crosslinked (pink rhombus) to an Acrydite-modified oligonucleotide adapter (pink). This crosslinking occurs through a photo-crosslinking reaction between ^CNV^K and pyrimidines via a [2+2] cycloaddition upon UV-A irradiation (366 nm). Following the preparation of thick tissue slices (see Methods), the adapter (pink)-primer (black) complex and padlock (black) probes with unique gene identifiers (red) hybridize to intracellular mRNAs (gray dashed line) within the intact tissue. The probe set is copolymerized with acrylamide, forming a DNA-gel hybrid (blue wavy lines) through the adapter’s functionalized acrylic group, followed by the removal of unbound lipids and proteins. Subsequently, enzymatic ligation and rolling circle amplification (RCA) construct *in situ* cDNA amplicons. These cDNA amplicons are further anchored into the hydrogel network via hydrogel re-embedding. Barcodes on the unique gene identifiers are read out via cyclic *in situ* sequencing with error reduction by dynamic annealing and ligation (SEDAL). This comprehensive quantification of RNA enables the elucidation of gene expression patterns and the identification of cell types within the native 3D tissue context. **c**, Left: Schematics and representative fluorescent images of negative and positive control experiments in 100 µm tissue sections of the mouse cerebral cortex. Using a 5′ Acrydite adapter photocrosslinked with the primer produces equivalent results to direct 5′ Acrydite modification of the primer, both surpassing the performance of adapter-primer hybridization alone or hydrogel physical retention. Right: Quantification of cell images showing the average amplicon reads per cell (n=4 images per condition). Red: DNA amplicons from 4 cell type markers. Blue: DAPI. Scale bar: 20 µm. Two-sided independent *t*-test, ****P < 0.0001. Data shown as mean ± standard deviation. **d**, Left: Schematics and representative fluorescent tissue images of 6 rounds of sequencing with and without cDNA re-embedding. In the absence of cDNA re-embedding, PEGylated bis(sulfosuccinimidyl)suberate (BSPEG) is used to crosslink cDNA. This results in background accumulation and reduced cDNA detection efficiency. Fluorescent images show Ch1 to Ch4 (color-coded channels for barcode decoding) and cell nuclei (blue) in mouse brain slices. Right: Quantification of cell images showing the average amplicon retention rate after 6 rounds of sequencing (n=4 images per condition). Two-sided independent *t*-test, ****P < 0.0001. Data shown as mean ± standard deviation. **e**, Representative raw fluorescent tissue images across 200 µm and quantification of DNA amplicon signal intensity at different tissue depths. **f**, Deep-RIBOmap probe design: Primer (black) and padlock (black) probes with unique gene identifiers (red) hybridize to intracellular mRNAs (gray dashed line), while splint probes (green) bind to the 18S rRNA of ribosomes. Splint probes serve as splints for proximity ligation and circularization of padlock probes. Both the primer and splint probe feature a flanking linker sequence at their 5′ ends and are covalently crosslinked (pink rhombus) to an acrydite-modified oligonucleotide adapter (pink).

STARmap employs a padlock probe, designed to target specific mRNA species of interest, along with a primer that binds to the same mRNA transcript adjacent to the padlock probe binding site. In the previous publication of the STARmap protocol adapted for thick tissue^30^, each primer contains a 5′ Acrydite modification to anchor the probe into the hydrogel. However, this modification is expensive to synthesize and not scalable for large gene numbers. In this report, Deep-STARmap incorporates a common “flanking linker sequence” at the 5′ end of all primers. This addition enables an Acrydite-modified adapter to hybridize and covalently crosslink with the flanking linker, allowing the whole probe set of primers to be conjugated to polyacrylamide hydrogels during polymerization (**Fig. 1b**). Covalent crosslinking is achieved with a nucleoside analog, 3-cyanovinylcarbazole nucleoside (^CNV^K)^41^, incorporated into the adapter. Upon 366 nm UV irradiation, the ^CNV^K-containing adapter undergoes rapid photocrosslinking to the complementary strand via an adjacent pyrimidine base, a process shown to be non-damaging to DNA^42,43^. Experimental optimization revealed that an adapter-to-primer ratio of 5:1 is sufficient for complete conversion of primer, and higher ratios do not increase the number of amplicons in mouse brain samples (Extended Data Fig. 1a-c). Notably, probes with a photocrosslinked 5′ Acrydite adapter performed equivalently to those with 5′ Acrydite modifications incorporated during solid phase synthesis (**Fig. 1c**). It is worth noting that the photocrosslinked adapter approach is markedly more efficient and scalable, as it employs a universal flanking linker sequence and corresponding adapter for all primers to allow pooled synthesis. In contrast, attachment of 5′ Acrydite for each individual probe during solid phase synthesis is extremely costly, especially in the setting of >5,000 probes with up to 70 bases in length. Due to its high multiplexing capability, our method enables the embedding of a large number of probe sets into the hydrogel, expanding the number of targetable RNA species from dozens to thousands. Our findings demonstrate that UV crosslinking significantly enhances probe incorporation efficiency, resulting in a higher detection yield of cDNA amplicons compared to mere adapter-primer hybridization. This method substantially outperforms the approach of relying solely on hydrogel physical retention (**Fig. 1c**). We also demonstrated that anchoring probe sets into the hydrogel is more efficient than the previously reported strategies^40,44^ of anchoring RNA molecules into the hydrogel in our experimental setting. (Extended Data Fig. 1f, g).

After hydrogel polymerization using a mixture of redox initiator and thermal initiator to embed the tissue and polymerizable primers (Extended Data Fig. 1d, e), we performed protein digestion, enzymatic ligation, and rolling circle amplification (RCA) to construct *in situ* cDNA amplicons. We observed that primer polymerization alone could not efficiently retain cDNA amplicons as puncta for more than 4 imaging cycles, potentially because they are prone to displacement, disassemble, and even fragmentation caused by buffer-dependent hydrogel expansion and contraction between imaging cycles, resulting in progressively lower SNR (**Fig. 1d**). To maintain the position and integrity of the amplicons through multiple detection cycles, a second round of hydrogel embedding was introduced following RCA, which outperformed several alternative re-embedding strategies (Extended Data Fig. 2a-d). Collectively, the implementation of these strategies devised explicitly for Deep-STARmap significantly enhances its robustness and scalability, enabling consistent spatial transcriptomics readouts across 200-µm thick sections of the mouse brain (**Fig. 1e**).

Next, we leveraged the insights gained from developing Deep-STARmap to establish Deep-RIBOmap for investigating spatial translatomics in thick tissue samples. RIBOmap utilizes a tri-probe design strategy to selectively detect and amplify ribosome-bound mRNAs: in addition to the padlock and primer, an additional splint DNA probe hybridizes to ribosomal RNAs (rRNAs)^45^. Building upon this design, Deep-RIBOmap incorporates a “flanking linker sequence” at the 5′ end of both the primer and the splint DNA probe (**Fig. 1f**). An Acrydite-modified adapter covalently crosslinks to these flanking linkers, enabling the integration of the entire tri-probe set into polyacrylamide hydrogels during polymerization. Using the same workflow as Deep-STARmap, Deep-RIBOmap achieves spatial translatomic profiling in thick tissue blocks.

### Deep-STARmap and Deep-RIBOmap in mouse brain with 1,017 genes

To evaluate the scalability of Deep-STARmap and Deep-RIBOmap for high-throughput 3D intact tissue transcriptomic and translatomic sequencing, we applied these techniques to thick mouse brain sections (Methods), targeting a curated list of 1,017 genes. This gene list was compiled from reported cell-type marker genes in adult mouse CNS single-cell RNA sequencing (scRNA-seq) datasets and spatial transcriptomic mouse brain atlases^46–49^. Gene identities encoded by five-nucleotide sequences on the padlock probes were read out through six rounds of sequencing by ligation with error rejection (SEDAL).

We performed pairwise Deep-STARmap and Deep-RIBOmap mapping on adjacent 150-µm-thick coronal sections of the mouse hemisphere, encompassing multiple brain regions (198,675 cells for Deep-STARmap and 164,029 cells for Deep-RIBOmap). To annotate cell types and align them with established nomenclature, we integrated our Deep-STARmap and Deep-RIBOmap with a published spatial brain atlas with curated cell typing annotations, using two different approaches independently. In the first approach, we used FuseMap^50^, a recently developed integration method that transfers cell type annotations leveraging both spatial and cellular information (**Figure 2a**). We also benchmarked the results using a second approach, where an established method, Harmony^51^, is used solely relying on single-cell gene expression information (**Figure 2b**). Both methods were applied independently to the same datasets and yielded consistent results: the confusion matrix of major cell type assignments showed that FuseMap’s cell types were highly concordant (82.4% matched labels) with those identified by the traditional single-cell sequencing integration method (**Fig. 2c**). Since FuseMap is a pre-trained model that integrates multiple large-scale spatial transcriptomic datasets and cell-type annotations of the mouse brain^50^ and has demonstrated higher accuracy in sublevel transferred annotations, we proceeded with FuseMap for downstream analyses.

**Fig. 2.**
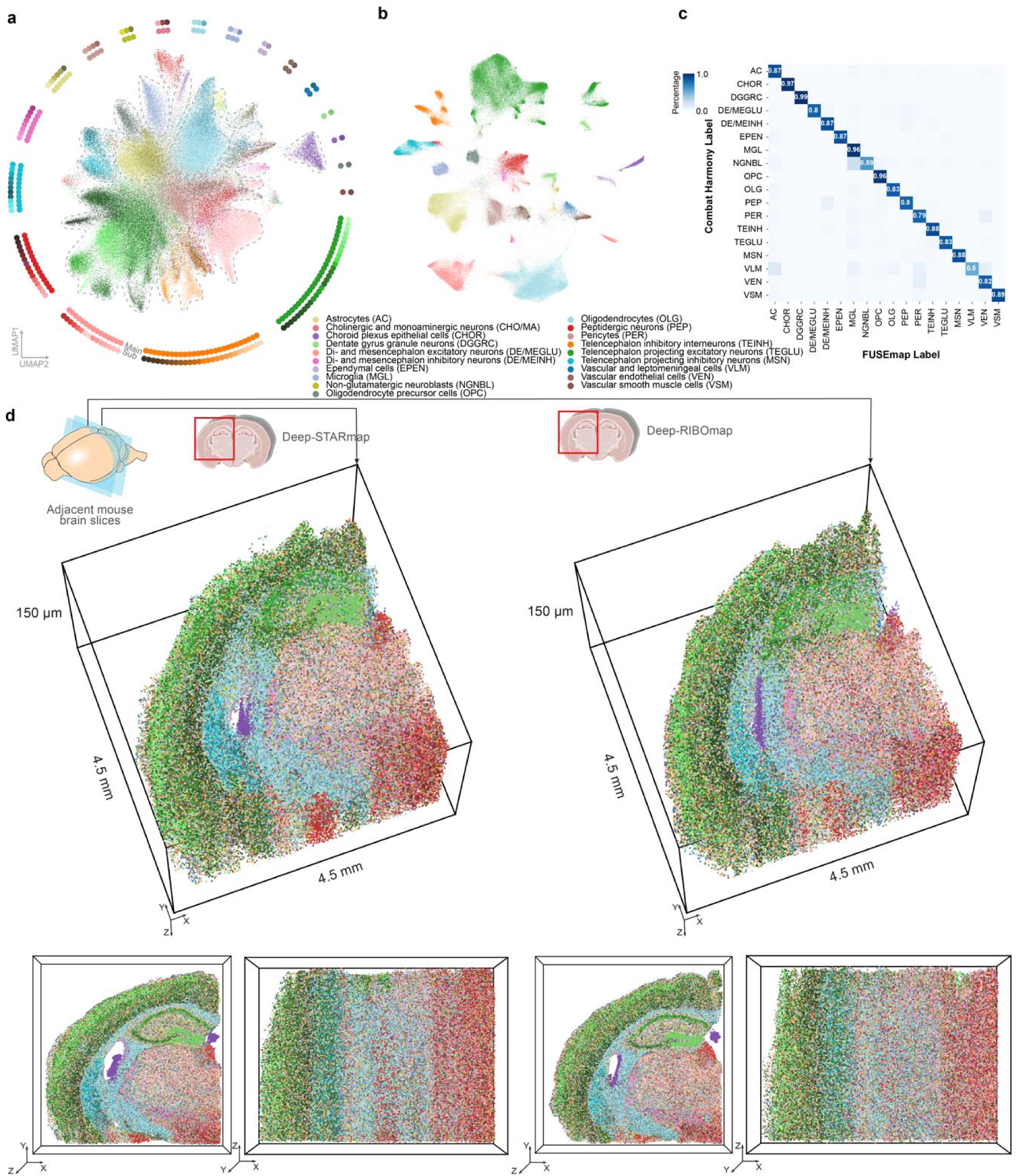
Spatial single-cell transcriptomic and translatomic profiling of 1,017 genes in thick mouse brain slices: **a-b**, Uniform Manifold Approximation and Projection (UMAP) plot visualizations of transcriptional and translational profiles of 362,704 cells collected from mouse coronal hemibrains using FuseMap (**a**) and integration using Harmony (**b**). Surrounding diagrams display 137 subclusters derived from 19 main clusters. **c**, Confusion matrix of cell type labels obtained from FuseMap and Harmony integration, visualizing cell types with more than 100 cells in the sample. **d**, 3D molecular cell-type maps derived from Deep-STARmap (left) and Deep-RIBOmap (right) across adjacent 150-µm thick sections from the mouse hemisphere. Each dot represents one cell, colored by its subcluster identity, using the same color code as in (a).

FuseMap integration, followed by nearest-neighbor label transfer, identified 19 main cell types, including 9 neuronal, 5 glial, 1 immune, and 4 vascular cell clusters, all of which exhibited canonical marker genes and expected spatial distributions. Further hierarchical clustering within each main cluster resulted in 137 subclusters (**Fig. 2d**). These major and subcluster annotations were consistent with previously published brain atlas datasets (**Fig. 2d**)^49,52,53^. Our spatial transcription and translation patterns of canonical cell-type marker genes and neurotransmitter genes aligned well with previously published spatial transcriptomic and translatomic sequencing results (Extended Data Fig. 3a, b). Based on these cell typing results, we generated spatial cell maps of the imaged hemibrain region. Our analysis demonstrated consistent cell typing between Deep-STARmap and Deep-RIBOmap in terms of gene expression patterns, cell-type composition, and spatial distribution of cell types (**Fig. 2d** and Extended Data Fig. 3c).

By exploiting the single-cell and spatial resolution of paired Deep-STARmap and Deep-RIBOmap datasets, we probed the heterogeneity in translational regulation across various cell types and brain regions. To investigate translationally regulated genes across different cell types, we first performed gene clustering using Deep-STARmap and Deep-RIBOmap profiles, identifying 18 gene modules (Extended Data Fig. 4a) with distinct functions and expression patterns (Extended Data Fig. 4b). Prior research has shown that non-neuronal cells, particularly oligodendrocytes, exhibit significant translational regulation^45^. We analyzed a gene module comprising 74 genes predominantly expressed across the oligodendrocyte lineage, from oligodendrocyte progenitor cells (OPCs) to mature subtypes (OLG1 and OLG2). Our findings recapitulate previous observations^45^, demonstrating that genes with higher translation efficiency in OPCs are linked to oligodendrocyte differentiation, while those with elevated translation efficiency in mature oligodendrocytes are associated with myelination (Extended Data Fig. 4c,). Beyond our extensive transcriptome analysis of the brain, we focused on investigating translational control at the subcellular level. Translation localized to the soma and processes in brain tissue plays a pivotal role in the organization and plasticity of neuronal and glial networks in response to physiological stimuli during neurodevelopment and memory formation (Extended Data Fig. 5a). To dissect this localized translation, we categorized Deep-RIBOmap reads in 3D thick tissue blocks into somata-localized reads (within the cell body, identified using Watershed 3D^54^) and processes-localized reads (the rest of the reads). We then identified the top 10% of genes with the highest and lowest processes-to-somata ratios, designating them as enriched in processes and somata, respectively (Extended Data Fig. 5b, d). Gene ontology (GO) analysis indicated that genes enriched in processes are involved in cell projection, cell junction, and cell-cell signaling, while those enriched in somata are associated with the extracellular matrix and various receptors (Extended Data Fig. 5c).

Given the ability of Deep-STARmap and Deep-RIBOmap to measure multiple layers of cells, we next tested whether our methods could resolve volumetric patterns of cell organization in 3D. We performed a detailed analysis of the nearest-neighbor distances among various interneuron subtypes. Prior studies have demonstrated that interneurons of identical subtypes frequently form juxtaposed pairs in the mouse visual cortex. Our result substantiates these findings, indicating that an inhibitory neuron is predominantly adjacent to another of the same subtype (Lamp5, Vip, Sst, or Pvalb) rather than other inhibitory subtypes (Extended Data Fig. 5e, f). This close spatial proximity might be related to the formation of gap junctions, which are crucial for synchronized firing patterns and may enhance visual responses in the cortex^55–57^.

### Single-cell morphology analysis of molecular cell types with Tetbow

Understanding the brain function necessitates a detailed mapping of its neuroanatomy. Electron microscopy (EM) remains the gold standard for neuroanatomical studies, offering nanometer-scale resolution^58,59^. However, EM reconstructions are largely incompatible with molecular cell-typing, resulting in a trade-off between spatial resolution and molecular information. Additionally, the current analytical throughput of EM is inadequate for studying the long-range spatial organization of mouse and mammalian neurons. The integration of stochastic multicolor labeling techniques^13,14,60^ with spatial transcriptomic mapping offers a promising solution. This combined approach enables the generation of comprehensive, co-profiling of transcriptome and morphology of individual neurons within densely labeled neural circuits.

To simultaneously interrogate transcriptomic readouts and morphology within single cells by exploiting the unique advantages of thick tissue mapping, we integrated the stochastic multicolor genetic labeling tool, Tetbow^14^, into our workflow (**Fig. 3a**). Tetbow enables bright and high-resolution mapping of intermingled neurons *in situ* by tagging individual neurons with stochastic combinations of three cytoplasmically-localized fluorescent proteins. It has also been demonstrated that systemically delivered AAVs allow a more uniform distribution of labeled cells and color diversity^61^. Thus, we utilized the AAV-PHP.eB^61^ variant to co-administer three separate vectors encoding three fluorescent proteins, along with a tTA expression vector to activate combinatorial fluorescent protein expression across the entire brain. (**Fig. 3b**, Extended Data Fig. 6a).

**Fig. 3.**
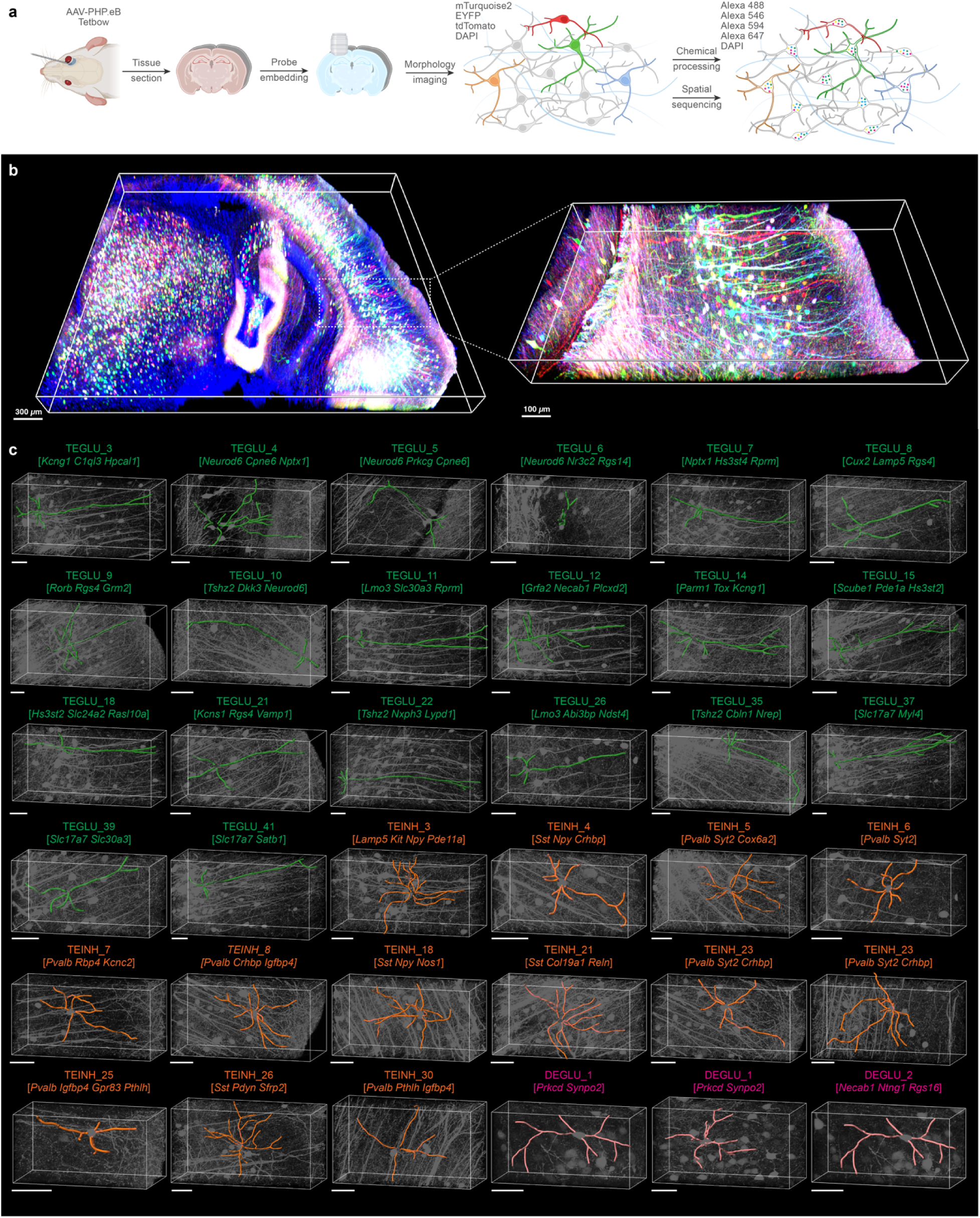
Characterizing the morphological features of transcriptomic types. **a**, Deep-STARmap combined with Tetbow enables simultaneous profiling of gene expression and neuron morphologies. AAV-PHP.eB delivers vectors encoding fluorescent proteins and the tTA expression vector. Following tissue sectioning and embedding probe sets into hydrogel, Tetbow fluorescent proteins and DAPI are imaged. After protein digestion, cDNA amplicons are constructed and sequenced. DAPI co-staining serves as a fiducial marker for image registration between FP images and *in situ* sequencing images, enabling the identification of Tetbow-labeled neurons by molecular subtype. **b**, Left: Volume rendering of neurons in the hippocampus and thalamus labeled with Tetbow. Neurons exhibit unique colors generated by the stochastic and combinatorial expression of three fluorescent proteins (tdTomato, EYFP, and mTurquoise2), enabling high-resolution identification and differentiation of individual neurons. Right: Zoom-in view of volume rendering of mouse cortical pyramidal neurons labeled with Tetbow. **c**, Representative individual morphological reconstructions of 30 transcriptome-defined subtypes of excitatory and inhibitory neurons. These reconstructions illustrate the distinct morphologies associated with each neuronal subtype, providing insights into the structural diversity within the neural network. Scale bar: 50 µm.

After tissue sectioning and embedding the probe sets into the hydrogel through polymerization, we performed imaging for the three Tetbow fluorescent proteins (FPs) along with DAPI, and observed bright, high-quality labeling of diverse neuronal cell types across all regions of the brain (**Fig. 3a**). Following morphology imaging, we digested the FPs from the sample using tissue clearing to enable subsequent transcriptome profiling of 1,017 genes. cDNA amplicons were constructed and sequenced as previously described. We additionally used DAPI as a fiducial marker for image registration between the two imaging modalities to correspond each FP-labeled neuron to its molecular subtype identity resolved by Deep-STARmap (Extended Data Fig. 6b, c).

To visualize the morphological diversity of labeled neurons, we established a semi-automated morphological reconstruction pipeline in Bitplane Imaris. In total, we reconstructed the dendritic arbors of 40 principal cells and interneurons within the imaged volume, spanning across 34 molecular subtypes (**Fig. 3c**). We focused our study on the dendritic arbor to maximize the accuracy of our traces, as it has been demonstrated that fine axonal morphologies cannot be consistently and faithfully recapitulate without a membrane-localized marker^60^. In agreement with past findings^62,63^, we were able to resolve the characteristic dendritic trees of principal pyramidal neurons in different brain regions, including CA1 hippocampal (TEGLU_3,7,8,9,10,11,12,14,15,18,21,22,26,35,37,39,41) and layer V neocortical pyramidal neurons (TEGLU_4,5,6). As expected, we resolved the most prominent dendritic structure of pyramidal neurons: the apical dendrite, which in CA1 hippocampal neurons extends towards the stratum lacunosum-moleculare (SLM), while in layer V neocortical neurons extend towards the cortical surface, both branching to form tree-like structures. Similarly, although cortical GABAergic inhibitory interneurons constitute a minority of the total neocortical neuronal population, we additionally confirmed that their morphology was resolvable with our approach, and that they exhibit a wide diversity of dendritic morphologies. In this study, we also elucidate the morphological diversity of several major subclasses of GABAergic neurons^64^, classified by their transcriptomic profiles, within the mouse cerebral cortex. In conclusion, we simultaneously profiled molecular cell types and morphologies at single-cell resolution in the adult mammalian brain in a scalable way.

### Deep-STARmap in human cutaneous squamous cell carcinoma

Deep-STARmap’s potential extends beyond neuroscience. One particularly promising application lies in the field of oncology, where a comprehensive understanding of the spatial organization of tumors, their microenvironments and immune interactions is crucial. Skin cancers account for ∼90% of all human malignancies. The second-most prevalent skin cancer is cutaneous squamous cell carcinoma (cSCC), which arises from keratinocytes (the major cell type of the epidermis). Over 1 million new cSCC cases are diagnosed annually in the United States^65^, with an estimated 3.7% of cSCCs leading to metastatic disease and 1.5% of cases resulting in death from disease^65^. The leading risk factor for cSCC is chronic ultraviolet radiation (UVR) exposure, which has mutagenic effects on the skin. UVR-induced somatic mutations translate to a large burden of tumor neoantigens that are thought to be responsible for the high immunogenicity of cSCCs^66^. Of note, immunosuppressed patients are at a 65-100 fold higher risk of developing cSCC and are significantly more likely to be diagnosed with multiple and metastatic cSCCs due to a failure of cancer immunosurveillance^67,68^. Immunotherapies such as immune checkpoint inhibitors have shown promise in the treatment of advanced cSCC^69^, however many patients fail to respond and the biomarkers, precise cell subpopulations, and mechanisms underlying response versus resistance are not well understood. There is great interest in assessing the spatial organization and signaling between tumor, immune, and stromal cells in the native tumor microenvironment. Prior spatial studies of cSCC have been limited to thin or 2D tissue samples that do not capture the full complexity of tumor architecture, as human skin’s barrier function makes it resistant to enzymatic digestion and macromolecule penetration. Thus, we applied Deep-STARmap to more comprehensively assess tumor organization and tumor-immune cell interactions in cSCC.

We curated a targeted list of 254 genes from previously published scRNA-seq studies of normal skin and skin cancers, including markers for common skin and immune cell types^70–73^. Deep-STARmap was performed on a 60-µm-thick section of human cSCC obtained from Mohs micrographic surgery (MMS), a sample that included both cSCC tumor and adjacent normal skin (Extended Data Fig. 7a). Following cell segmentation in 3D, data processing, and integration with a published cSCC scRNA-seq dataset^71^, we conducted cell typing and visualized cell clusters on the UMAP space based on single-cell RNA expression (**Fig. 4a**). 9 cell types were identified using known marker genes: keratinocytes, tumor-specific keratinocytes (TSKs), fibroblasts, endothelial cells, B cells, Langerhans cells, macrophages/dendritic cells (DCs), cytotoxic T cells, and regulatory T cells/exhausted T cells (**Fig. 4a, b**). Deep-STARmap enabled dissection of tumor spatial organization at single-cell resolution. Consistent with histologic tumor spatial patterns noted at the time of MMS, tumor-specific keratinocytes in this sample were primarily localized to the center of the tissue while non-tumor keratinocytes were localized to normal skin at the sample periphery (**Fig. 4c**).

**Fig. 4.**
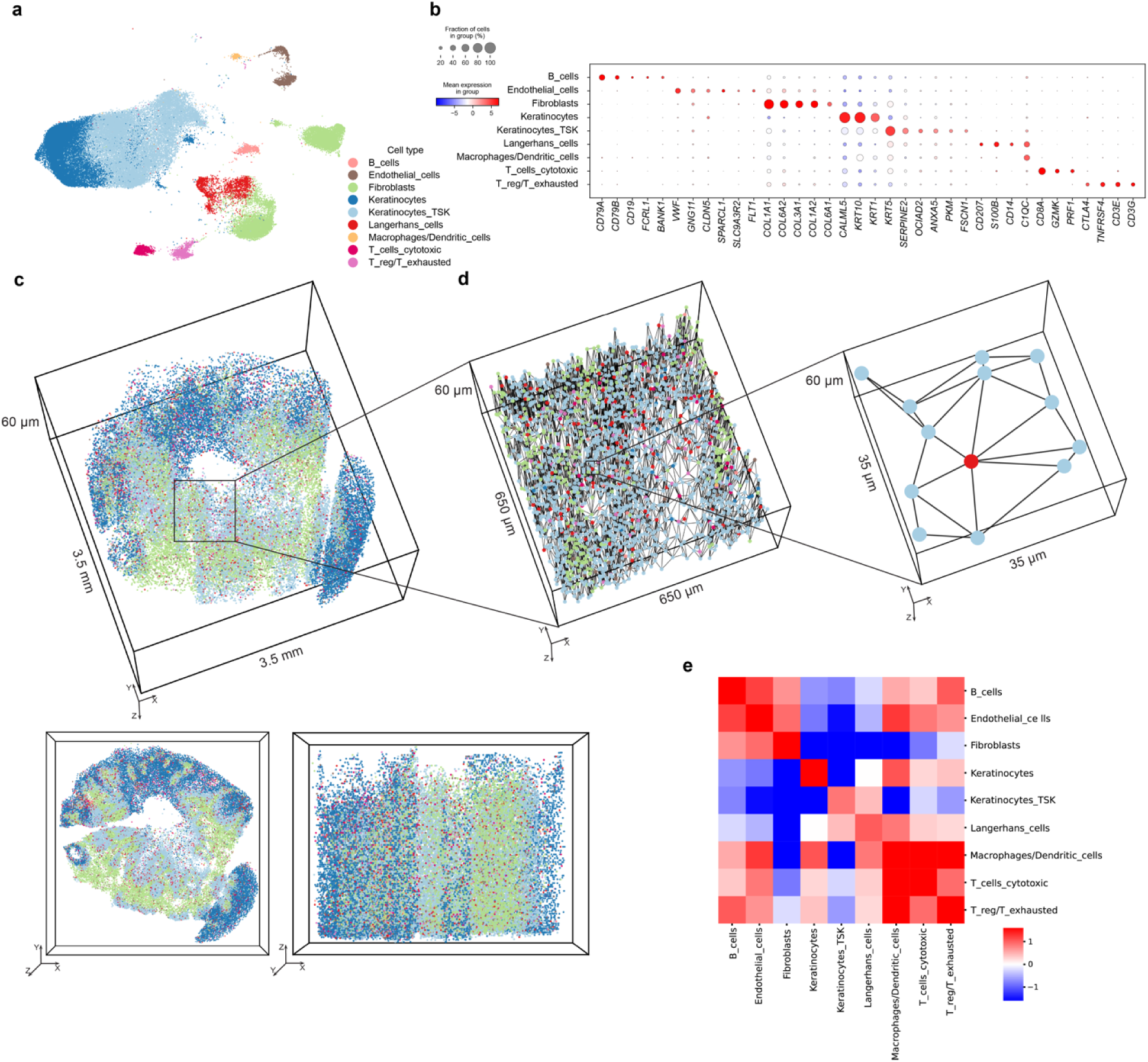
Mapping cell-cell interactions in human cSCC. **a**, UMAP plot visualization of transcriptional profiles of 51,471 cells, integrated using Harmony with a published cSCC scRNA-seq dataset. Cells are color-coded according to their cell-type identity. **b**, Dot plot illustrating the top differentially expressed marker genes for each major cluster. The color scale represents the log_2_ fold change in gene expression compared to the mean gene expression values across all cells. The dot size indicates the percentage of cells expressing the genes within each major cell type. **c**, 3D molecular cell-type maps generated from Deep-STARmap, using the same color coding as in (a). **d**, Zoomed-in view of the interaction between Langerhans cells and tumor-specific keratinocytes within a mesh graph of physically neighboring cells. Each cell is depicted as a spot colored according to its main cell type, with physically neighboring cells connected by edges. **e**, 3D cell-cell adjacency quantified by the normalized number of edges between pairs of cell types.

### Mapping cell-cell interactions in human cSCC

To characterize cell-cell interactions, we generated a mesh graph via Delaunay triangulation of cells and computed a near-range cell-cell adjacency matrix from spatial connectivity as previously described^74,75^. This allowed us to identify the nearest neighbors of each cell and to quantify the number of edges between cells of each type with cells of the same or other cell types. A heat map of cell type frequencies among first-tier neighbors revealed clear patterns of cell type-specific cell-cell communication (**Fig. 4d**). The same analysis was performed on a thin 15-µm section of the cSCC sample taken within the same 3D volume (Extended Data Fig. 7b, c). As expected, more cell-cell contacts were detected in thick tissue (mean of 14.3 connected cells) compared to thin tissue (mean of 6.0 connected cells) (Extended Data Fig. 7d, e).

Across the cSCC sample, strong interactions were detected among cells of the same type, with more same cell type interactions identified in thick tissue compared to pseudo-thin tissue (Extended Data Fig. 7b, c). Similarly, immune cell interactions with other immune cell types such as macrophages/DCs with T cells were more strongly detected in thick compared to pseudo-thin tissue (Extended Data Fig. 7b, c). Interestingly, tumor-specific keratinocytes only interacted strongly with two cell types: other tumor-specific keratinocytes or Langerhans cells (**Fig. 4d, e**). This was again more evident in thick tissue than thin tissue, demonstrating that the additional 3D morphological information provided by Deep-STARmap increases the sensitivity and robustness for quantifying cell-cell contacts.

Langerhans cells (LCs) are the major resident antigen-presenting cells of the skin and are known to interact with keratinocytes via E-cadherin. LCs have been reported to encounter cSCC cells prior to other DC subtypes^76^ and stimulate cytotoxic CD8 T cells and NK cells more efficiently than other DC subsets^77^. In our cSCC sample, LCs interacted with T cells and tumor-specific keratinocytes, but not normal keratinocytes outside the tumor, indicating tumor-specific immune responses (**Fig. 4e**). Taken together, our Deep-STARmap cSCC data identified a disease-relevant interaction between tumor-specific keratinocytes and Langerhans with more accurate and quantitative spatial distribution compared to thin tissue analyses.

## Discussion

In this study, we present Deep-STARmap and Deep-RIBOmap as novel imaging platforms for *in situ* transcriptomic and translatomic sequencing within intact tissue blocks. To enable robust performance and scalability over existing approaches, we introduced new strategies for thick tissue RNA imaging, including scalable probe synthesis, efficient probe anchoring, and robust cDNA crosslinking. These technological developments are pivotal for scaling up 3D *in situ* transcriptomic and translatomic profiling to encompass thousands of genes and across larger tissue regions. This scalability facilitates the integration of molecular characterizations with morphology mapping in neuroscience within thick tissue blocks. We demonstrated that Deep-STARmap and Deep-RIBOmap could profile the transcription and translation of over 1,000 genes within intact thick mouse brain tissue sections, significantly expanding the readouts from larger cell populations. Incorporating combinatorial fluorescence labeling using the Tetbow system allowed high-throughput *in situ* co-profiling of spatial transcriptomics and single-neuron morphology in thick tissue blocks, enabling multimodal mapping on a volumetric scale previously unattainable. For example, our platform is potentially compatible with MAPseq^15^ and BARseq^16^ to uncover the organizing principles of neuronal circuitry in thick tissue blocks. Moreover, our platform can be further applied to decode the spatial transcriptomics and translatomics of specific neurons with activity dynamics being collected by live imaging.

Our platform can also be generalized to study various heterogeneous cell populations in diverse tissues. We demonstrated that our 3D *in situ* profiling platform is adaptable for profiling difficult-to-digest human skin cancer samples, providing more accurate and quantitative measurements of tumor-immune spatial patterns. Furthermore, we anticipate that our 3D *in situ* profiling platform will be highly useful for studying human organoid cultures, which are extensively used to replicate *in vivo* 3D organ development from 2D embryonic germ layers during organogenesis^78^. These organoids, typically measuring hundreds of micrometers, necessitate *in situ* profiling in both healthy and diseased states to advance our understanding of human tissue development, pathology, and therapeutic responses.

In summary, 3D *in-situ* spatial transcriptomics and translatomics, exemplified by Deep-STARmap and Deep-RIBOmap, offer a robust methodology for integrating molecular data with high-resolution cellular imaging. This comprehensive approach allows for detailed analysis of anatomical and functional dynamics within tissues. Such techniques are poised to substantially enhance our understanding of the underlying mechanisms of tissue functionality and pathology, thereby facilitating deeper scientific exploration and potential therapeutic innovations.

## Author contributions

X.S. and X.W. conceived the idea and developed Deep-STARmap and Deep-RIBOmap for the study. X.S. carried out experimental work, performed *in situ* sequencing, and conducted computational and data analyses. J.A.L. designed the gene lists for human cSCC, acquired samples, and made significant contributions to the analysis of human cSCC. S.L., Y.H., and Z.T. performed data analysis. Z.L. and J.L. helped with method optimization. Y.Z. helped with AAV packaging and conducted animal work. W.X.W helped with morphology analysis. X.S., J.A.L., and X.W. wrote the manuscript with input from all authors. X.W. supervised the study.

## Acknowledgments

The authors thank B.E.Deverman lab for helping with AAV packages. The authors thank H.Shi, A.Roy, F.Kostas, S.Furniss for their helpful feedback and comments during manuscript writing. X.W. gratefully acknowledges support from Edward Scolnick Professorship, gift from Stanley Center for Psychiatric Research, Escaping Velocity Award, Ono Pharma Breakthrough Science Initiative Award, Merkin Institute Fellowship, and NIH DP2 New Innovator Award (1DP2GM146245), and NIH BRAIN CONNECTS (UM1 NS132173). J.L. acknowledges support from NIH/NIDDK 1DP1DK130673. J.A.L gratefully acknowledges support from the NIH T32 Dermatology Training Program Grant (T32AR007098-47) and Dermatology Foundation Dermatologist Investigator Research Fellowship. W.X.W is a Damon Runyon–National Mah Jongg League, Inc. Fellow, supported by the Damon Runyon Cancer Research Foundation (DRG#: 2512-23).

## Methods

### Mouse lines

All animal procedures adhered to the care guidelines approved by the Institutional Animal Care and Use Committee (IACUC) of the Broad Institute of MIT and Harvard, under animal protocol #0255-08-19. For the experiments, C57/BL6 mice aged between 6 to 10 weeks were procured from The Jackson Laboratory (JAX). Mouse were housed with 4∼5 animals per cage with arbitrary food and water in a room with 18∼23 °C temperature and 40–60% humidity under a 12-h light-dark cycle.

### Human cutaneous squamous cell carcinoma samples

Human cutaneous squamous cell carcinoma tissue was obtained from deidentified discarded hospital specimens approved under the Massachusetts General Hospital Research Committee/IRB protocol #2013P000093.

### Tetbow AAV injections

The AAV plasmids utilized in this study include pAAV-TRE-mTurquoise2-WPRE (Addgene #104110), pAAV-TRE-EYFP-WPRE (Addgene #104111), pAAV-TRE-tdTomato-WPRE (Addgene #104112), and pAAV-ihSyn1-tTA (Addgene #99120). Tetbow components were packaged into AAV.PHP.eB as previously described^61^. In brief, for each capsid, HEK 293T cells (ATCC CRL-3216) were transfected with a combination of pAAV plasmid and two AAV packaging plasmids (kiCAP-AAV-PHP.eB and pHelper) in a 1:4:2 weight ratio, using polyethylenimine, with a total of 40 μg of DNA per 150-mm culture dish. Fluorescence expression, when applicable, was evaluated via microscopy, and the media was refreshed 20-24 hours post-transfection. Viral particles were collected 72 hours post-transfection from both the cells and the medium by centrifugation, forming cell pellets. These cell pellets were then resuspended in a buffer containing 500 mM NaCl, 40 mM Tris, 10 mM MgCl_2_, pH ∼10 and 100 U/mL of salt-activated nuclease (SAN, 25 U/μL, Arcticzymes, 70910-202) and incubated at 37°C for 1.5 hour. Following incubation, the cell lysates were subjected to centrifugation at 2,000g to remove cellular debris. The viral particles were then isolated through a series of iodixanol gradient steps (15%, 25%, 40%, and 60%). Viruses were collected from both the 40/60% interface and the 40% iodixanol layer. The concentration of the viral particles and buffer change were achieved using Pierce™ Protein Concentrators (Thermo Scientific, 88528), and they were subsequently suspended in sterile phosphate-buffered saline (PBS). To quantify viral titers, viral genomes were measured using quantitative PCR (qPCR). The procedure included treating samples with DNase I (Roche Diagnostics, 4716728001) to eliminate non-packaged DNA and subsequently with proteinase K (Roche Diagnostics, 03115828001) to digest the viral capsid, thereby exposing the viral genomes for qPCR quantification. A linearized genome plasmid served as the reference standard. The viral titers for tTA, tdTomato, EYFP, and mTurquoise2 were 2.15 × 10^13^, 2.31 × 10^13^, 3.04 × 10^13^, and 2.63 × 10^13^ vg/ml, respectively.

Intravenous administration of AAV.PHP.eB mixture (1 × 10^11^ vg tTA, 3.33 × 10^11^ vg tdTomato, 3.33 × 10^11^ vg EYFP, 3.33 × 10^11^ vg mTurquoise2) was performed via injection into the retro-orbital sinus of adult female C57BL/6 mice (8-10 weeks of age). Twenty-eight days post-injection, the mice were anesthetized with isoflurane. Transcardial perfusion was carried out, initially with 50 mL of cold PBS, followed by 50 mL of 4% PFA. The entire brain was then post-fixed in 4% PFA at 4°C for 3 hours. Subsequently, the brain was washed multiple times with PBS and placed in a 30% sucrose solution (in PBS) at 4°C overnight or until it had sunk. Finally, the brain was embedded in O.C.T. (Fisher, 23-730-571) and frozen in liquid nitrogen and stored at −80 °C. Thick tissue sections were prepared and carefully transferred into pretreated glass-bottom plates.

### Chemicals and enzymes

Chemicals and enzymes listed as name (supplier, catalog number): 12-Well Plate, No. 1.5 Coverslip, 14 mm Glass Diameter, Uncoated (MatTek, P12G-1.5-14-F); PlusOne Bind-Silane (Sigma, 17-1330-01); 16% PFA, EM grade (Electron Microscope Sciences, 15710-S); Methanol (Sigma-Aldrich, 34860-1L-R); Tween-20, 10% solution (Teknova, T0710); Triton-X100, 10% solution (Sigma-Aldrich, 93443-100ML); 10X PBS (Thermo Fisher, 70011044); 1X PBS (Thermo Fisher, 10010049); 20X SSC buffer (Thermo Fisher, 15557044); Methacrylic acid N-hydroxysuccinimide ester, 98% (Sigma-Aldrich, 730300-1G); Acrylamide solution, 40% (Bio-Rad, 161-0140); Bis Solution, 2% (Bio-Rad, 161-0142); Ammonium persulfate (Sigma-Aldrich, A3678-100G); N,N,N′,N′-Tetramethylethylenediamine (Sigma-Aldrich, T9281-50ML); OmniPur SDS, 20% (Calbiochem/Sigma, 7990-200ML); NeuroTrace Fluorescent Nissl Stains, yellow (Molecular Probes/Fisher Scientific, N21480); COVER GLASS CIRCLE, 12mm, #2, 1oz/BX (Electron Microscopy Sciences, 72226-01); Gel Slick Solution (Lonza, 50640); Formamide, Deionized (Sigma aldrich, 4650-500ML); Antarctic Phosphatase Reaction Buffer (New England Biolabs, B0289S); Antarctic Phosphatase (New England Biolabs, M0289L); BSA (New England Biolabs, B9200S); Glycine (Sigma aldrich, 50046-250G); Ribonucleoside Vanadyl Complex (New England Biolabs, S1402S); DMSO, anhydrous (Invitrogen/Thermo Fisher, D12345); DNase/RNase-Free Distilled Water (Invitrogen/Thermo Fisher, 10977023); 4’,6-diamidino-2-phenylindole (DAPI) (Thermo Fisher, 62248); Acetic acid (Sigma-Aldrich, A6283-100ML); Poly-D-Lysine (Thermo Fisher, A3890401); dNTP mix (thermofisher, 18427089); 5-(3-aminoallyl)-dUTP (Invitrogen, AM8439); BSPEG9 (thermofisher, 21582); Proteinase K Solution (Invitrogen, 25530049); SUPERase•In RNase Inhibitor (Thermo Fisher, AM2696); T4 DNA Ligase (Thermo Fisher, EL0012); Phi29 DNA Polymerase (Thermo Fisher, EP0094); Yeast tRNA (Thermo Fisher, AM7119)

### Deep-STARmap and Deep-RIBOmap probe design

The Deep-STARmap and Deep-RIBOmap padlock and primer probes were developed based on the methodologies outlined in Wang et al. and Zeng et al., with specific modifications^30,45^. Each Deep-STARmap and Deep-RIBOmap primer incorporated a “flanking linker sequence” (CCTACCAGTACGACGTATTTAGCAA) at the 5′ end to enable hybridization with an Acrydite-modified oligonucleotide. The Deep-RIBOmap additionally required a splint probe, composed of three segments: a 25-nucleotide sequence at the 5′ end complementary to the 18S ribosomal RNA (rRNA), a stretch of 50 deoxyadenosine nucleotides (dA), and a 12-nucleotide padlock template at the 3′ end. To prevent the 3′ terminus of the splint probes from serving as an RCA primer, a 3′ Inverted dT modification was included. Additionally, each splint probe incorporated a “flanking linker sequence” (CCTACCAGTACGACGTATTTAGCAA) at the 5′ end to facilitate the hybridization process with the Acrydite-modified oligonucleotide.

### Adapter and primer pre-treatment

The ^CNV^K-containing adapter ([5Acryd]GCTA[^cnv^K]ATACGTCGTACTGGTAGG[Inv-dT], ordered from Gene Link with PAGE purification) undergoes rapid photo cross-linking to the complementary strand through an adjacent pyrimidine base upon UV irradiation. The irradiation process was conducted using the Boekel UV Crosslinker (234100) equipped with 368 nm-wavelength bulbs (Boekel Part Number 920-0307). The adapter to primer was maintained at a molar ratio of 5:1.

### Deep-STARmap and Deep-RIBOmap protocol

Glass-bottom 12-well plates (Mattek, P12G-1.5-14-F) were treated with oxygen plasma using the Anatech Barrel Plasma System at 100W and 40% O_2_ for 5 min. Following this, the plates were immersed in a 1% methacryloxypropyltrimethoxysilane (Bind-Silane) solution for 60 min at room temperature. The plates then underwent three consecutive ethanol washes and were allowed to air dry. Subsequently, a 0.1 mg/mL Poly-D-lysine solution was applied to the plates for 1 hour, followed by three rinses with distilled water.

Tissue slices were transferred and adhered to pre-treated glass-bottom 12-well plates. The samples were permeabilized using 1 mL of pre-chilled methanol at −20°C for one hour. During this period, PBST solution, comprising 0.1% Triton-X 100 in PBS, was prepared. The samples were then washed with 500 µL of PBSTR (0.1 U/mL SUPERase·In in PBST) for 30 min. This was followed by a quenching step with 500 µL of quenching solution (1 mg/mL Yeast tRNA, 100 mM Glycine in PBSTR) at room temperature for 30 minutes, followed by another 30-min wash with PBSTR. Subsequently, hybridization buffers were prepared. The base composition of the hybridization buffer included 2× SSC, 10% formamide, 1% Triton-X 100, 20 mM RVC, 0.1 mg/mL yeast tRNA, 0.1 U/µL SUPERase·In, and 0.2% SDS. For the Deep-STARmap samples, this buffer was supplemented with pooled Deep-STARmap padlock and pre-treated primer at a concentration of 5 nM per oligo. For the Deep-RIBOmap samples, the hybridization buffer additionally contained 100 nM of pre-treated splint probe for RIBOmap. The samples were incubated in 300 µL of hybridization buffer in a 40°C humidified oven with gentle shaking for 36 hours. After incubation, the samples were washed for 30 min with PBSTR, followed by a 30-min wash in high salt buffer (4× SSC in PBSTR) at 37°C. Finally, the samples were washed once more with PBSTR at 37°C.

To cast the tissue-hydrogel hybrid, the samples were first incubated with monomer buffer (4% acrylamide, 0.2% bis-acrylamide, 2× SSC) supplemented with 0.2% TEMED and 0.25% VA-044 at 4°C for 60 min. Following incubation, the buffer was aspirated, and 55 µL of a polymerization mixture (0.2% TEMED, 0.2% ammonium persulfate, and 0.25% VA-044 in monomer buffer) was added to the center of the sample and immediately covered with a Gel Slick-coated coverslip. The polymerization process was conducted in a 40°C N_2_ oven for 90 min. Subsequently, the sample was washed with PBSTR three times for 15 min each. For Tetbow samples, the tissue was stained with DAPI for 3 hours and then immersed in a washing and imaging buffer (10% formamide in 2× SSC buffer) containing 0.1 U/µL SUPERase·In RNase inhibitor. Confocal images of Tetbow fluorescent proteins (tdTomato, EYFP, and mTurquoise2) and DAPI were acquired using an inverted confocal microscope, Leica TCS SP8 (version 3.5.5.19976), equipped with a 405 nm and 442 nm diode, a white light laser, HyD detectors, and a 25× water-immersion objective (NA 0.95). The voxel size for imaging was 0.32 μm × 0.32 μm × 0.70 μm. The following wavelengths were used for imaging: 405 nm for DAPI, 442 nm for mTurquoise2, 506 nm for EYFP, and 550 nm for tdTomato.

The tissue-gel hybrids were then digested with 1 mL Proteinase K mixture (0.4 mg/mL Proteinase K in 2× SSC and 1% SDS) at 37°C for overnight, then washed by PBSTR 3 times for 30 min each. The sample was then incubated in ligation mixture (0.25 U/µL T4 DNA ligase, 1:100 BSA, 0.2 U/µL SUPERase·In RNase inhibitor) at room temperature overnight with gentle shaking and then washed with PBSTR three times for 30 mins each. Then the sample was incubated with 400 μl rolling-circle amplification mixture (0.5 U/µL Phi29 DNA polymerase, 250 µM dNTP, 20 µM 5-(3-aminoallyl)-dUTP, 1:100 BSA and 0.2U/µL of SUPERase·In RNase inhibitor in 1X Phi29 buffer) at 4°C for 60 min for equilibrium before incubating at 30 °C for 8-14 hours for amplification and then washed with PBST 3 times for 30 mins each. The samples were then treated with 20 mM methacrylic acid N-hydroxysuccinimide ester in 100 mM sodium bicarbonate buffer for 4 hours to overnight at room temperature. Following the exact same procedures casting tissue-hydrogel hybrid, cDNA amplicons were re-embedded with 2% acrylamide, 0.05% bis-acrylamide to enable cDNA amplicon crosslinking in the tissue-hydrogel setting, and such cross-linking is essential to maintain the position and integrity of the amplicons through many cycles of detection. Samples were stored in PBST or wash and imaging buffer at 4°C until imaging and sequencing.

Before SEDAL, the samples were treated with the dephosphorylation mixture (0.25 U/μL Antarctic Phosphatase, 1× BSA, in 1× Antarctic Phosphatase buffer) at 37 °C for 4 hours and washed by PBST three times for 30 min each. Each sequencing cycle began with treating the sample three times, 15 min each, with the stripping buffer (60% formamide and 0.1% Triton X-100 in water) at room temperature, followed by washing with PBST three times for 15 min each. Then the samples were incubated with a at minimal 300 µL sequencing-by-ligation mixture (0.2 U/μL T4 DNA ligase, 1× BSA, 10 μM reading probe, and 5 μM fluorescent decoding oligonucleotides in 1× T4 DNA ligase buffer) at room temperature for overnight, followed by rinsing with washing and imaging buffer three times for 10 min each before imaging. Images were acquired using the same Leica TCS SP8 with a 25× water-immersion objective (NA 0.95). The voxel size for imaging was 0.32 μm × 0.32 μm × 0.70 μm. For each round, images were acquired with Alexa 488, 546, 594, and 647 illumination. DAPI was dissolved in wash and imaging buffer and used for nuclei staining for 3 hours before the first round. The DAPI signal was collected at the first cycle of imaging with an additional 405 nm wavelength. Six cycles of imaging were performed to detect 1017 genes.

### Data processing for Deep-STARmap and Deep-RIBOmap

#### Deconvolution

Image deconvolution was achieved with Huygens Essential version 23.4.0 (Scientific Volume Imaging, The Netherlands, http://svi.nl). We applied the classic maximum likelihood estimation method with a signal-to-noise ratio of 10 and 10 iterations.

#### Image registration, spot calling, and barcode filtering

For image registration, spot calling, and barcode filtering, we utilized our custom software package, Starfinder (https://github.com/wanglab-broad/starfinder). This software corrects chromatic aberrations, enhances signals, registers images, and extracts positive reads (amplicons). Adjustments were made to accommodate the large datasets generated by thick tissue profiling. In short, image clarity is enhanced by intensity normalization and histogram equalization where images in the first sequencing round are used as reference. To ensure accurate and reliable identification of each cDNA amplicon’s barcode, we utilized a two-step registration process. First, we conducted a global registration using 3D fast Fourier transform. Next, we applied a non-rigid registration using MATLAB v.2023b’s ‘imregdemons’ function. This method adjusts for any shifts and distortions between imaging sessions, ensuring precise alignment of the same amplicon’s positions across different sequencing rounds. Since the amplicon size is larger than amplicons in thin tissue, we applied a medium filter with ‘*medfilt2’* function in a 3-by-3 or 2-by-2 (depending on the average amplicon size) neighborhood around the corresponding pixel in the input image. Dots with intensity at their centroids less than the threshold were removed. The process of identifying individual amplicons in 3D was carried out using the ‘imregionalmax’ function in MATLAB to find local maxima within the images from the first sequencing round. The dominant color for each amplicon across all rounds of sequencing was then determined by estimating the amplicon size and integrating the voxel volume intensity in each channel. Each dot’s color composition was represented by an L2-normalized vector with four elements, and dots showing multiple maximum values within this vector were excluded. Initial filtering of dots was based on quality scores, which were computed as the average of –log(color vector value in the dominant channel) across all sequencing rounds. This metric quantified the degree to which each dot in each sequencing round was derived from a single color rather than a blend of colors. Subsequently, the barcode codebook was translated into color space, following the expected color sequence of the two-base encoded barcode DNA sequence. Only dots that met the quality threshold and had a matching barcode sequence in the codebook were retained, with all others being discarded. The 3D physical locations and gene identities of these filtered dots were then preserved for subsequent analysis.

#### 3D segmentation

3D image segmentation was performed based on the DAPI staining image and the composite image containing amplicon channels to create reference segmentations as previously described with minor adjustments^45,74,79^. Unlike thin tissue analysis, where images are stitched before segmentation, this approach is impractical for thick tissue profiling because the stitched files are too large for effective segmentation. Therefore, segmentation was performed on each field of view (FOV) individually, and the identified amplicons were stitched afterward. For each FOV, images targeting different cellular compartments were first processed using a median filter and then binarized with an automatically determined threshold in FIJI. Distance Transformed Watershed 3D was subsequently applied to generate a 3D segmentation mask for each cellular region. Connected components (objects) with fewer than 500 voxels were removed from the binary image. Finally, the images were dilated using a disk structure element with a radius of 10.

#### Reads assignment and stitching

Filtered amplicons overlapping each segmented cell region in 3D were assigned to their respective regions to compute a per-cell gene expression matrix. The TileConfiguration file generated from FIJI grid stitching was then used to merge detected amplicon signals from each FOV, ensuring the removal of duplicated cells and associated reads. Further strategies to exclude low-quality cells were applied as previously described in thin tissue analysis^30,45,74^.

#### Cell type classification via FuseMap

Cell type classification was performed using transfer learning with a pretrained FuseMap model, as previously described^50^. This model maps and annotates new query data with cell-type labels based on cell embeddings. In this study, a previously published brain spatial atlas served as the reference for training the FuseMap model, while thick tissue sections were used as the query datasets for annotation.

#### Harmony integration

To benchmark FuseMap performance, Harmony integration was employed. First, Deep-STARmap data were combined with Deep-RIBOmap data after preprocessing, followed by batch correction using the pp.combat function. Harmony integration was then applied to the combined dataset to create a joint PCA embedding^80^. A k-nearest neighbor (KNN) classifier was trained on the integrated PC space using cosine distance as the metric. This classifier was used in a label transfer process to annotate each cell based on its neighboring reference cells in the KNN graph. The label transfer was performed for the annotation at the “Rank4_Refine” level.

### Gene Ontology (GO) enrichment analysis

GO enrichment analysis was conducted using the DAVID database (https://david.ncifcrf.gov/)^81,82^. gProfiler (https://biit.cs.ut.ee/gprofiler/gost) was utilized for GO analysis. Enriched GO terms were selected from biological processes (BP) and cellular components (CC) with FDR < 0.05 for both cell-type-resolved Deep-STARmap and Deep-RIBOmap profiles, as well as for somata-enriched translation genes and processes-enriched translation genes.

### Gene Clustering

The gene expression (*log2_norm1e4*) of the 4 samples were first averaged across the cell types within each sample, respectively. Subsequently, the average expression values were standardized by calculating the Z-score within each sample. The standardized vectors were merged and clustered with the Leiden algorithm from Scanpy^83^ (Version 1.9.3).

### Near-range cell–cell adjacency analysis

Near-range cell-cell adjacency analysis was performed to quantify the number of edges between cells of each main cell type and cells of other main cell types, as previously described^75,84^. The adjacency value between cell types A and B was defined as the number of A-B edges within a 1-hop neighborhood on the Delaunay tissue graph, calculated using scipy.spatial. Raw counts were normalized against a null distribution created by 1,000 random spatial shifts of cells.

### Morphological reconstructions

3D reconstructions of single-neuron morphologies were generated from 3D image stacks using Imaris (Oxford Instruments; v. 9.7.2-10.1.1). Dendritic arbors of Tetbow-labeled neurons were initially reconstructed semi-automatically with the filament tracer in autodepth mode. These reconstructions were then extensively manually corrected and curated using the filament tracer in manual mode. A fully connected neuronal structure was reconstructed wherever possible while remaining faithful to the image data. Any processes that could not be definitively linked to the main structure were left unconnected.

---

**Extended Data Fig. 1.**
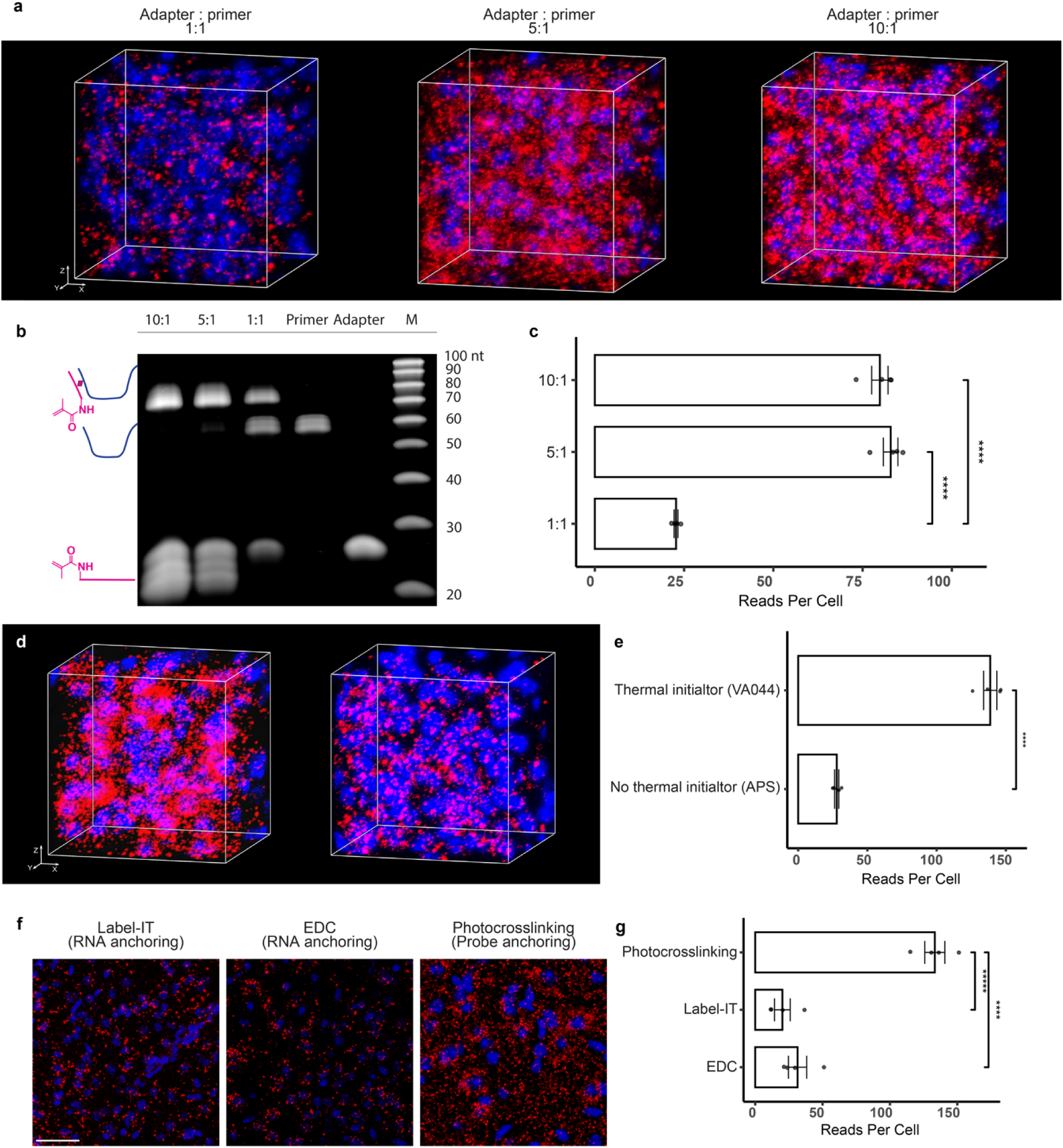
Optimization of probe crosslinking. **a**, Representative fluorescent imaging illustrating probe anchoring efficiency in a hydrogel matrix with various adapter-primer ratios. Red: DNA amplicons from 4 cell type markers. Blue: DAPI. Scale bar: 10 µm. **b**, 15% TBE-Urea gels demonstrating UV crosslinking efficiency with varying adapter-primer molar ratios. ^CNV^K- and Acrydite-containing adapter used for UV crosslinking is [5Acryd]GCTA[cnvK]ATACGTCGTACTGGTAGG[Inv-dT] (24 nt). Primer used is 58 bp ssDNA with a 24 nt flanking liner at the 5′ end. M, Marker: IDT ssDNA 20/100 Ladder. **c**, Quantification of cell images showing the average amplicon reads per cell (*n*=4 images per condition). Two-sided independent *t*-test, \*\*\*\**P* < 0.0001. Data presented as mean ± standard deviation. **d**, Representative fluorescent imaging demonstrating probe anchoring efficiency with and without the use of the VA-044 thermal initiator in the first round of polymerization. Red: DNA amplicons from 4 cell type markers. Blue: DAPI. Scale bar: 10 µm. **e**, Quantification of cell images showing the average amplicon reads per cell (*n*=4 images per condition). Two-sided independent *t*-test, \*\*\*\**P* < 0.0001. Data presented as mean ± standard deviation. **f**. Representative fluorescent imaging demonstrating detection efficiency of covalently anchored RNA molecules or probes within the hydrogel in the Deep-STARmap setting. Red: DNA amplicons from 4 cell type markers. Blue: DAPI. Scale bar: 50 µm. **g**, Quantification of cell images showing the average amplicon reads per cell (*n*=4 images per condition). Two-sided independent *t*-test, \*\*\*\**P* < 0.0001. Data presented as mean ± standard deviation.

**Extended Data Fig. 2.**
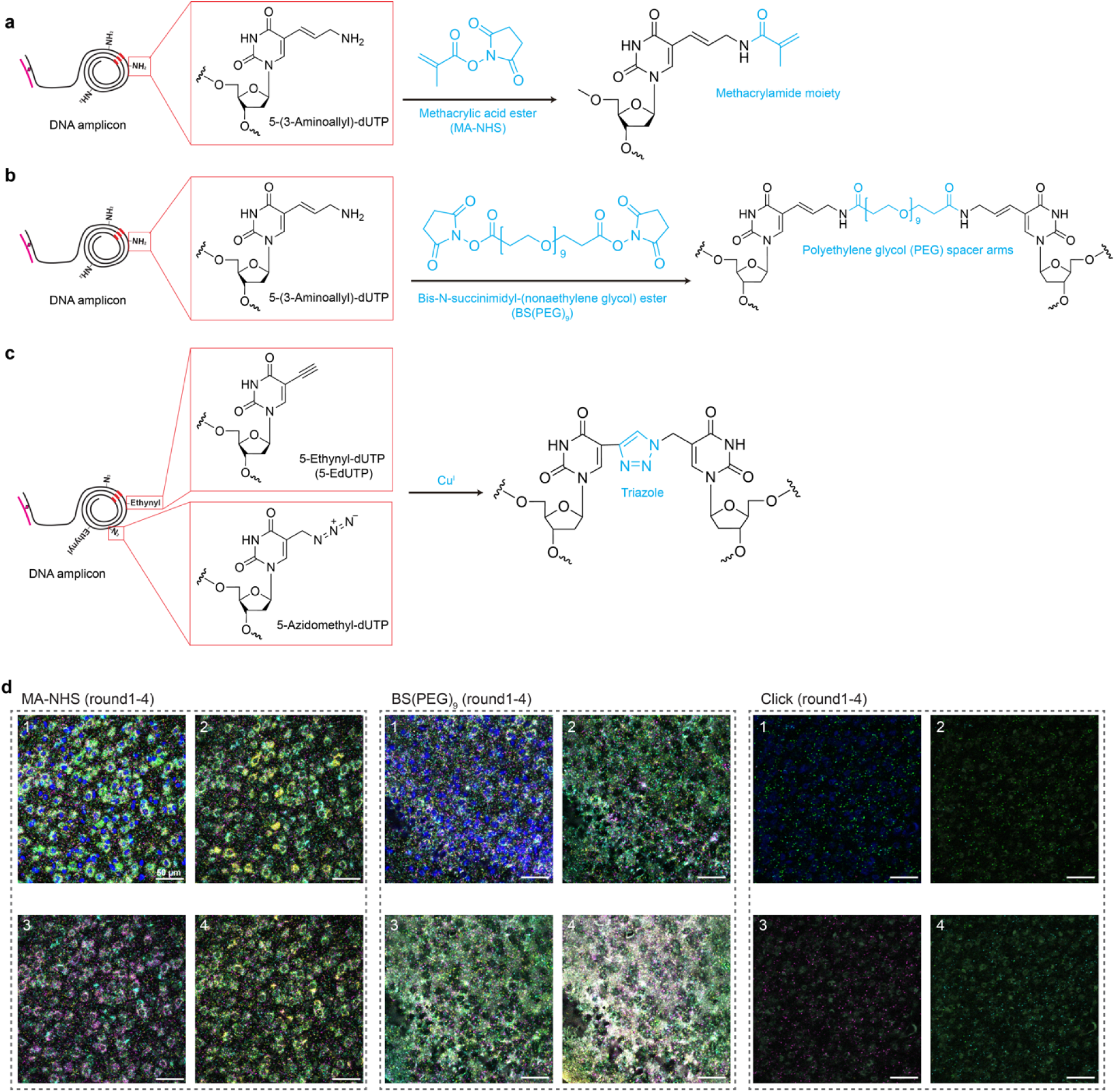
Optimization of re-embedding strategy. **a**, Mechanism of cDNA crosslinking using hydrogel re-embedding. Amine-modified nucleotides were incorporated into the RCA reaction. MA-NHS enables rapid conjugation to nucleophilic groups on the amplicons via its NHS ester under mild conditions. These functionalized methacrylamide moieties are then integrated into the hydrogel, effectively immobilizing the cDNA amplicons. **b**, Mechanism of cDNA crosslinking using BSPEG. Amine-modified nucleotides were incorporated into the RCA reaction followed by BSPEG crosslinking, where the NHS esters of BSPEG react with the amino groups on the amplicons. **c**, Mechanism of cDNA crosslinking using Click chemistry. Azide and alkyne groups were incorporated during the RCA process, followed by the addition of copper to catalyze the azide-alkyne cycloaddition, forming a stable triazole ring as a crosslinking method. **d**, Representative fluorescent imaging demonstrating sequencing signal-to-noise ratio using different cDNA crosslinking strategies. BSPEG and Click chemistry crosslinking result in higher background noise compared to hydrogel re-embedding after several rounds of sequencing. Additionally, the incorporation of azide and alkyne moieties during RCA significantly reduced amplification efficiency, leading to fewer amplicons.

**Extended Data Fig. 3.**
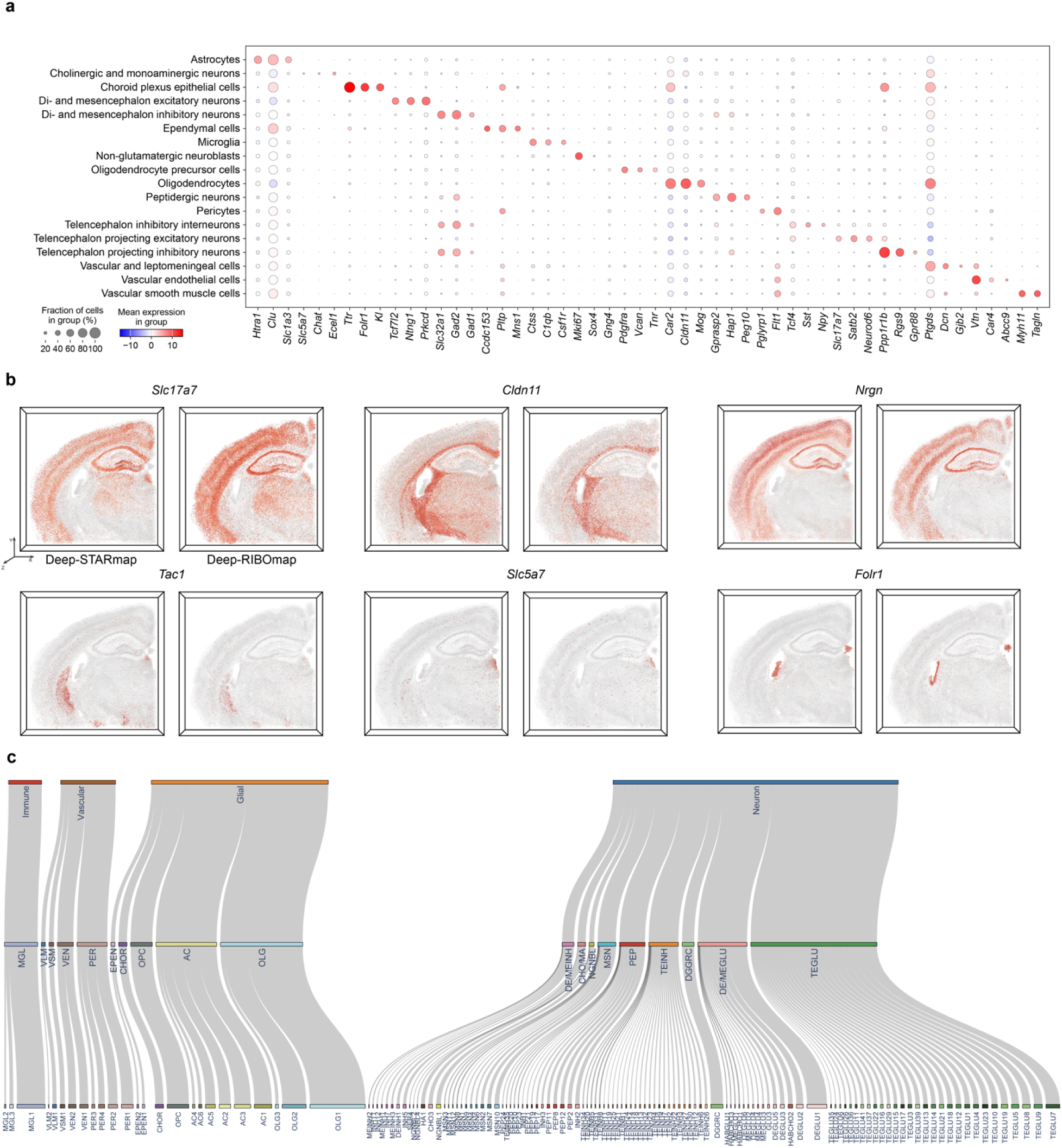
Spatially single-cell transcriptomic and translatomic profiling of 1017 genes in the mouse brain. **a**, Dot plot illustrating the expression levels of representative markers across various major cell types using Deep-STARmap and Deep-RIBOmap. The color scale represents the log_2_ fold change in gene expression compared to the mean gene expression values across all cells. The dot size indicates the percentage of cells expressing the genes within each major cell type. xyz size: 4.5 mm, 4.5 mm, 150 µm. **b**, Deep-STARmap (left) and Deep-RIBOmap (right) images of example cell marker genes and neurotransmitter genes. **c**, Hierarchical taxonomy of cell types showing the main level and subtype level cell-type identification and annotations.

**Extended Data Fig. 4.**
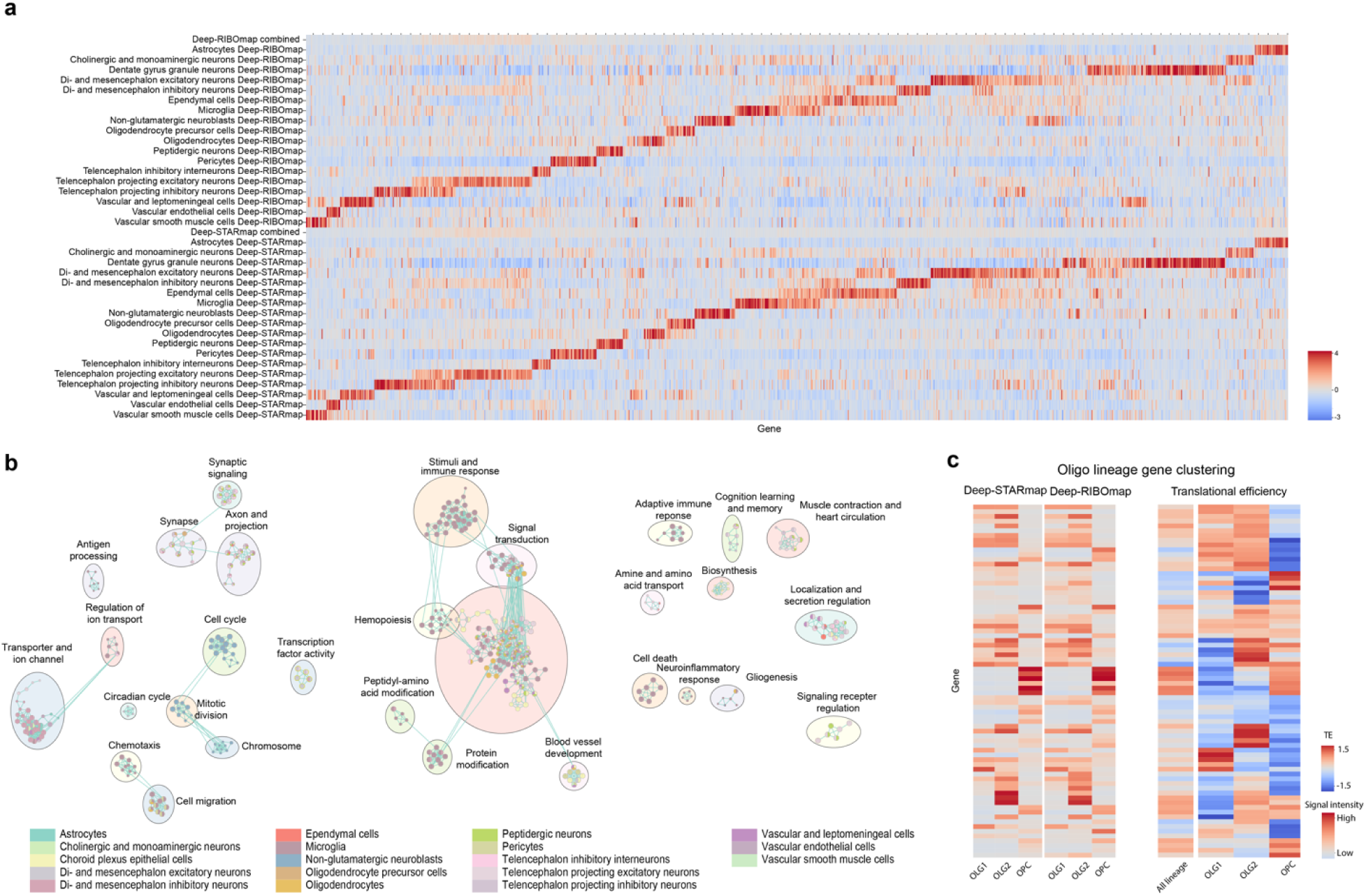
Comparison of spatial translatome and transcriptome in the mouse brain and cell-cell adjacency analysis. **a**, Heatmap showing the gene clustering using the RIBOmap and STARmap results by cell type (Z-score expression). **b**, Visualization of enriched GO terms within each gene module, categorized and color-coded by module. In the enrichment map, nodes represent enriched GO terms, with the size of each node reflecting the number of genes associated with that term. Edges between nodes indicate shared genes among the GO terms. **c**, Heatmap displaying gene clustering based on Deep-STARmap and Deep-RIBOmap results across the three oligodendrocyte lineage cell types (left). The right panel shows the translational efficiency (TE) of these genes within each oligodendrocyte lineage cell type (Z-score expression).

**Extended Data Fig. 5.**
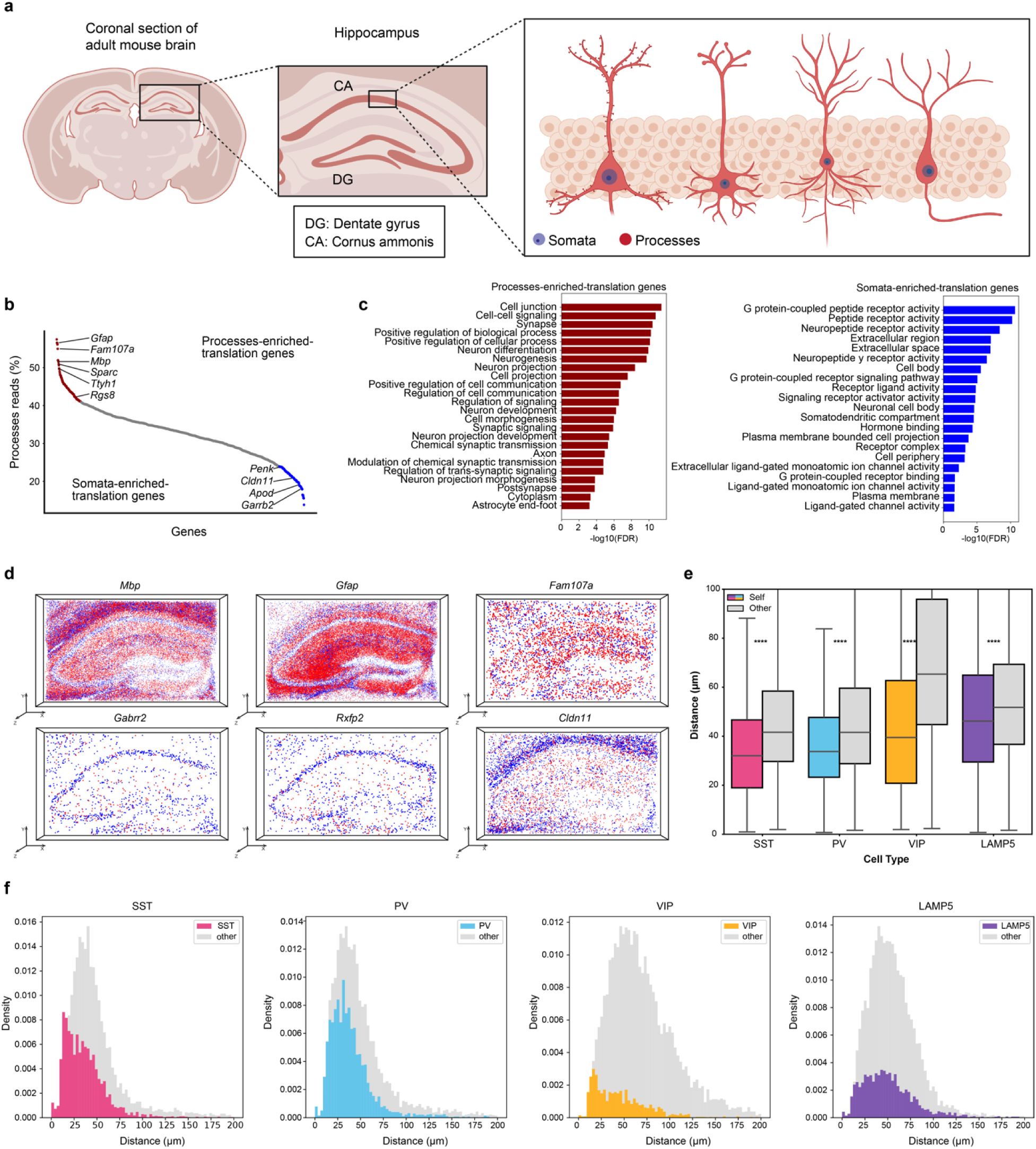
Localized translation in the somata and processes of neuronal and glial cells in the mouse brain. **a**, Schematic illustration of a hippocampal slice highlighting the somata and processes of hippocampal neurons. **b**, Processes read percentages of individual translating genes with genes rank-ordered based on their processes reads percentage. **c**, Significantly enriched GO terms for processes-enriched and somata-enriched translating genes. **d**, Spatial translation map of representative genes with enriched translation in processes (top) and somata (bottom) within the hippocampus, depicting somata reads in blue and process reads in red. **e-f**, Nearest-neighbor distance distributions in Deep-STARmap sample, comparing distances from cells in specific inhibitory neuronal subclasses to cells within the same subclass (“to self”) and to cells in different subclasses (“to other”).

**Extended Data Fig. 6.**
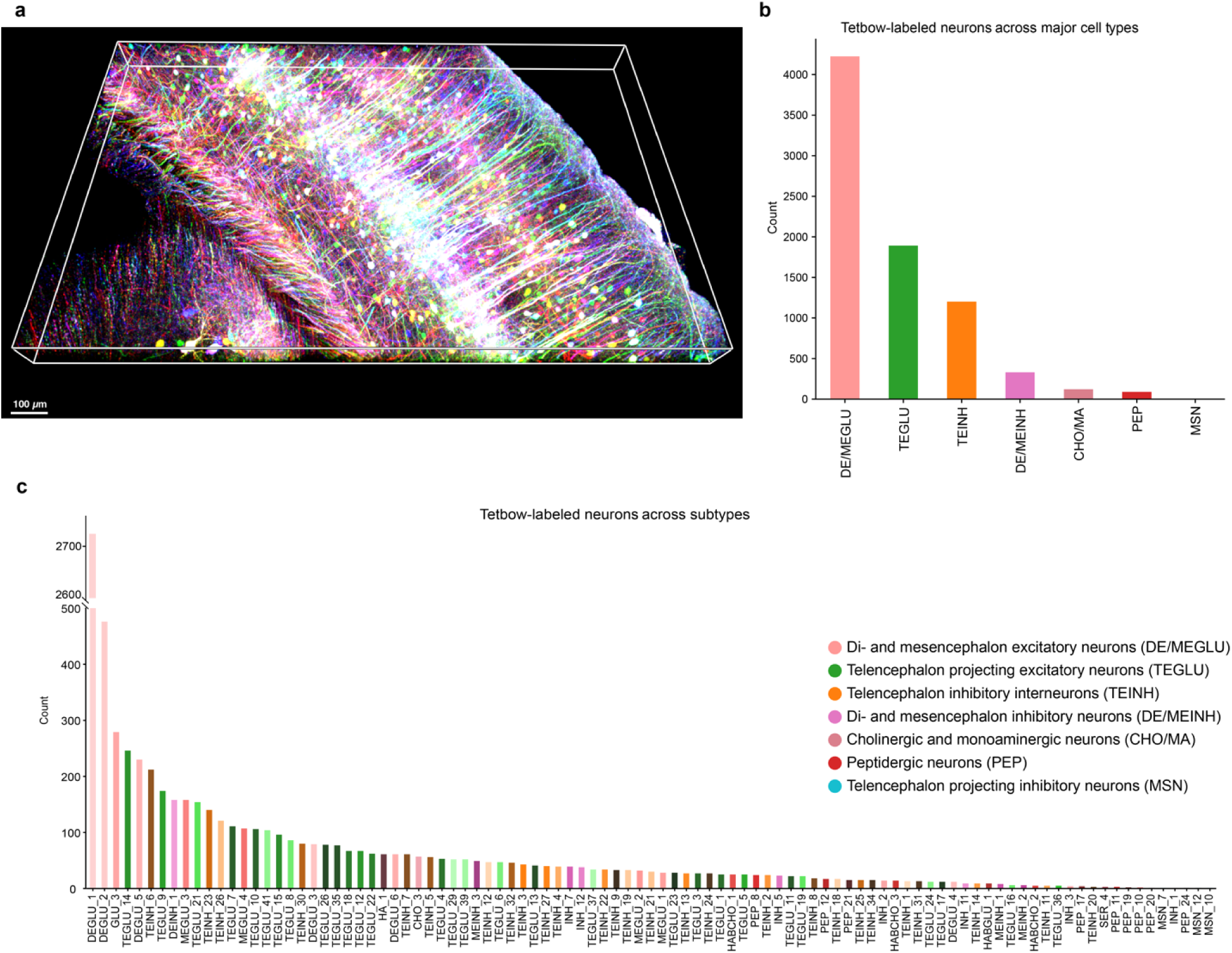
Quantification of Tebow-labeled neurons. (**a**) Another zoom-in view of volume rendering of mouse cortical pyramidal neurons labeled with Tetbow. **(b,c)** Cell count quantification of Tebow-labeled neurons across major cell types (b) and subtypes (c).

**Extended Data Fig. 7.**
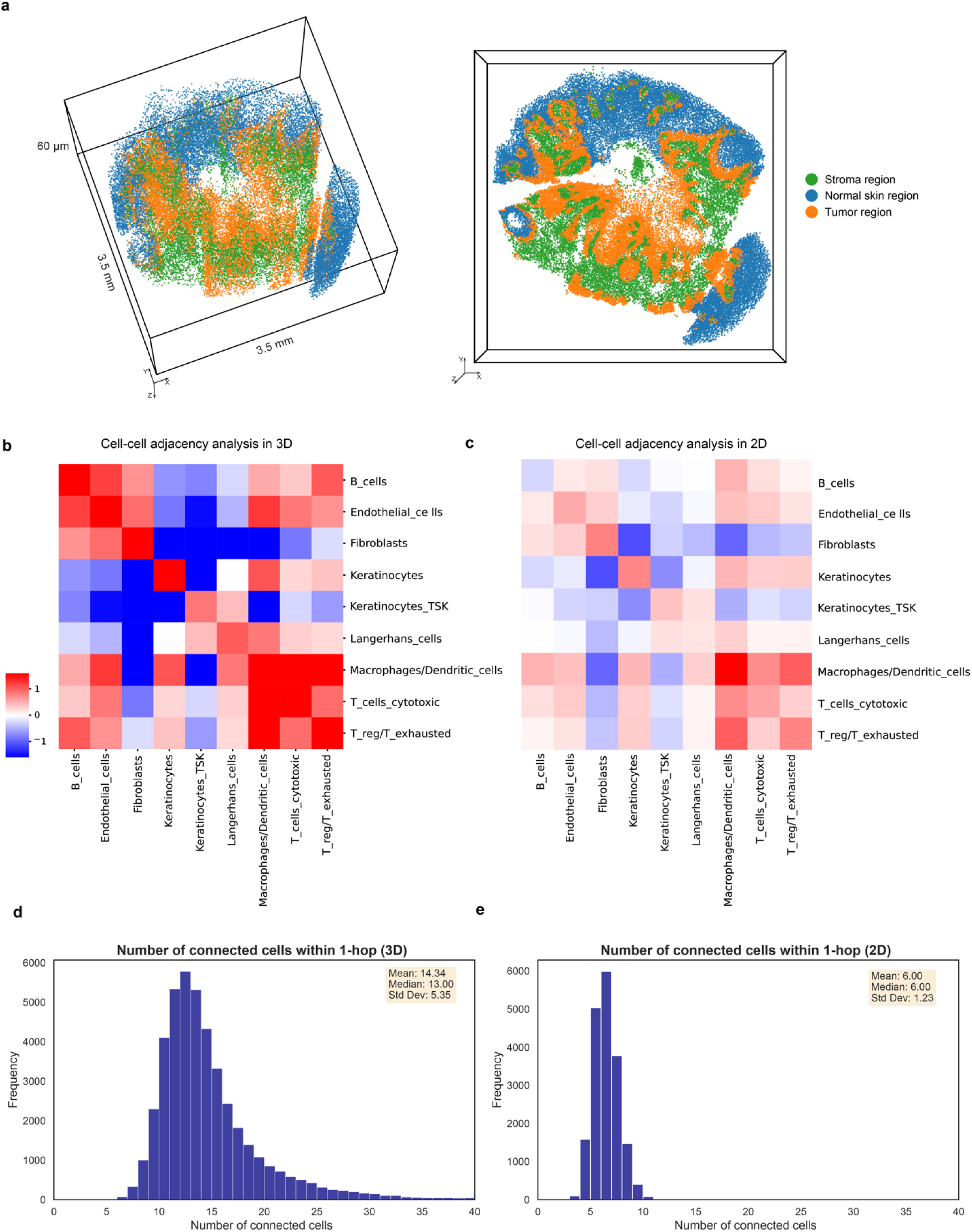
Cell-cell adjacency analysis in 2D and 3D. **a**, Molecular tissue region maps visualized in 3D. Each dot represents a cell. Three molecular regions can be identified: tumor region, fibroblast region, and normal skin region. These regions were identified by analyzing low-frequency, large-scale patterns within the spatial neighbors graph. **b-c**, Quantification of cell-cell adjacency in 3D (b) and 2D (c) by the normalized number of edges between pairs of cell types. The 2D analysis is performed by projecting 15 μm (∼1 cell layer) slices along the z-axis, taken within the same 3D volume as shown in Fig. 4. The 3D analysis reveals stronger cell-cell adjacency enrichment. **d-e**, The 3D analysis detects stronger cell-cell interactions because the number of connected cells (edges of a given cell in the mesh graph via Delaunay triangulation) is greater than in 2D. The 2D nearest-neighbor distances cannot accurately represent the 3D cellular environment.

